# Early threat perception is independent of later cognitive and behavioral control. A virtual reality-EEG-ECG study

**DOI:** 10.1101/2023.02.22.529523

**Authors:** Juanzhi Lu, Selma K. Kemmerer, Lars Riecke, Beatrice de Gelder

**Affiliations:** Department of Cognitive Neuroscience, Faculty of Psychology and Neuroscience, Maastricht University, Maastricht, Limburg 6200 MD, The Netherlands

**Keywords:** Control success, Electroencephalography, Social threat, Virtual Reality

## Abstract

Research on social threat has shown influences of various factors, such as agent characteristics, proximity and social interaction on social threat perception. An important, yet understudied aspect of threat experience concerns the ability to exert control over the thread. In this study, we used a Virtual Reality (VR) environment showing an approaching avatar that was either angry (threatening body expression) or neutral (neutral body expression) and informed participants to stop avatars from coming closer under five levels of control success (0, 25, 50, 75, or 100%) when they felt uncomfortable. Behavioral results revealed that social threat triggered faster reactions at a greater virtual distance from the participant than the neutral avatar. Event-related potentials (ERPs) revealed that the angry avatar elicited a larger N170/vertex positive potential (VPP) and a smaller N3 than the neutral avatar. The 100% control condition elicited a larger late positive potential (LPP) than the 75% control condition. In addition, we observed enhanced theta power and accelerated heart rate for the angry avatar vs. neutral avatar, suggesting that these measures index threat perception. Our results indicate that perception of social threat takes place in early to middle cortical processing stages, and control ability is associated with cognitive evaluation in middle to late stages.

## 1. Introduction

The ability to detect threat and react adaptively is a major evolutionary endowment of many species (LeDoux & Daw, 2018). Human and non-human studies of defensive behavior have documented different kinds of behavior in the face of threat, mainly freezing and fleeing (Eilam, 2005). Freezing has been defined as a threat-anticipatory state whereby an individual is hyperattentive to an environmental, potentially threatening signal, presumably also enhancing its processing (Blanchard et al., 1986; Livermore et al., 2021; Mobbs & Kim, 2015; Terburg et al., 2018). Previous work has investigated freezing-like reactivity using threat-related social stimuli, such as facial expressions and affective films (Hagenaars et al., 2014; Roelofs et al., 2010; Stins et al., 2011), as well as computer-based tasks (e.g., a gun shooting task) (Gladwin et al., 2016) and most recently whole body expression (de Borst & de Gelder, 2022; de Gelder et al., 2010; Mello et al., 2022). Bradycardia, a reduction in one’s heart rate, and reduced postural mobility are two principal physiological components of the freezing state in the face of threats (Roelofs et al., 2010). This pattern of physiological and behavioral activation is especially coordinated by the subcortical connections between amygdala nuclei - the basolateral nucleus receiving multisensory information and the central nucleus sending the main projections out - and the periaqueductal gray, the hypothalamus, and the rostral ventrolateral medulla (George et al., 2019). Stimulation of this circuit activates the sympathetic and parasympathetic nervous systems, which in turn coordinate switches from passive defensive states (e.g., freezing) to active defensive behavior (e.g., flight or fight) (Livermore et al., 2021; Terburg et al., 2018).

A critical factor for freezing-like reactions in humans is the proximity of the threat. Studies on peripersonal space (PPS), the proximate space surrounding the body where interactions with environmental stimuli occur (Bufacchi & Iannetti, 2018; Di Pellegrino & Làdavas, 2015; Serino, 2019), have shown that personal distance is an important determinant of defensive behavior in social interactions (Bogdanova et al., 2021; Brozzoli et al., 2013; Cléry et al., 2015; Graziano & Cooke, 2006; Pellencin et al., 2018). The defensive reactivity to potentially threatening stimuli near the PPS is associated with reduced motor cortex excitability (Avenanti et al., 2012), increased physiological reactivity (Ruggiero et al., 2021), and enhanced neural processing of the target stimulus in brain regions involved in defensive behavior (Vieira et al., 2020). Moreover, the neural network underlying PPS has been shown to respond also to indicators of social threat, specifically in nearby space (de Borst et al., 2020; Ellena et al., 2021). A threatening character invading one’s personal space is associated with increased activity in ventral premotor cortex and intraparietal sulcus (areas that are part of the brain network coding PPS) as well as amygdala and anterior insula (de Borst & de Gelder, 2022). Another line of research using electroencephalography (EEG) has shown that threatening body expression impact early event-related potentials (ERPs), such as N170 and vertex positive potential (VPP) (Stekelenburg & de Gelder, 2004; Van Heijnsbergen et al., 2007). These electrophysiological measures of threat also interact with PPS. A behavioral ERP study used a modified version of a paper- and-pencil validated measure of comfortable interpersonal distance (CID) to explore how participants react to the threat of interpersonal distance invasion (Perry et al., 2013). Participants were instructed to imagine they were in the center of the room, and as a friend or stranger approached, they could press a key to show that they wanted to stop the person from coming closer. It was observed that the potential threat (approaching person) elicited larger N1 for strangers compared to friends, whereas friends/strangers had no significant effect on P1 and LPP components. These ERP responses occurred from 50 to 800ms. Besides distance, another critical factor for adaptive threat response is related to control over the threat. Threat experience may be reduced when, for example, threat escape or another behavioral control is possible (Terburg et al., 2018). Active control behavior refers to a sense of control that can reduce or stop the approaching threat (Iachini et al., 2016; Wendt et al., 2017).

A major obstacle in research on human behavior in the face of social threat is the difficulty of rendering threatening situations in a realistic manner and obtaining valid measures of human behavior and physiology. The use of virtual reality (VR) opens unique chances for this important research field (Monti & Aglioti, 2018; Parsons et al., 2017) as it allows participants to experience a threatening event in a controlled laboratory environment “as if” it was actually happening to them (de Borst & de Gelder, 2022; de Borst et al., 2020; Fusaro et al., 2016; Mello et al., 2022; Tieri et al., 2017). VR-based designs implementing social threat from avatars have successfully been used in behavioral, fMRI and EEG studies (de Borst & de Gelder, 2022; Mello et al., 2022; Stolz et al., 2019). Here, we combined VR with behavioral measures and measures of neural and cardiac activity to assess with millisecond temporal resolution the impact of the avatar emotion (angry/neutral) and various levels of threat-control success (0%, 25%, 50%, 75%, or 100%). Our goal was to measure how social threat is perceived under naturalistic conditions implemented in VR and whether the ability to effectively control the threat affects how the source of the threat is processed at behavioral, neural, and cardiac levels.

## 2. Methods

### 2.1 Participants

Thirty healthy right-handed participants were recruited for this study. All participants had normal or corrected-to-normal vision without brain injury, history of psychiatric disorder, or current psychotropic medication. Participants provided written consent at the beginning of the experiment. They earned 7.5EUR or received one credit point per hour of participation. Four participants’ data were rejected because they did not press a button in more than 50% of the trials. Twenty-six participants’ data were included in the analysis (13 females, 13 males; age range 18-29 years, mean = 24.65; standard deviation (SD) = 3.60). The Ethical Committee of Maastricht University approved the study, and all procedures conformed to the Declaration of Helsinki.

### 2.2 Design and procedures

#### 2.2.1 VR scenario

The VR scenario consisted of a dark and narrow urban street, in which an avatar expressing an angry or neutral emotion, appeared and subsequently approached the participant. The VR scenario was programmed in Unity 3D (Unity Technologies, US). The basic design of the angry (raised arms) and neutral (arms down) body expressions were similar to a previous study (Mello et al., 2022), while the VR environment and the task settings were new for the present study. Participants wore a VR headset (HTC VIVE) and explored the 3D VR world before the start of the experiment by walking along streets and moving their heads around to visually explore the surroundings. This served to make participants immersed in the VR environment.

#### 2.2.2 Experimental design

The VR environment and task were explained to the participants. Participants were told that an angry or emotionally neutral avatar would appear in a dark, urban environment and approach them. They were informed that pressing a control button (space bar) could stop the avatar from coming closer, and they were encouraged to do so as soon as they felt uncomfortable. At the beginning of each trial, a cue appeared indicating the likelihood that pressing the space bar would effectively stop the avatar. There were five different controllable cue conditions (Fig1A). In the 0% condition, the button press never stopped the approaching avatar, while in the 100% condition pressing the space bar always stopped the avatar. In the intermediate conditions, the bar press stopped the avatar from approaching with 25%, 50% and 75% probability. A sketch of a trial is shown in Fig1B. All trials were preceded by a cue presented 1s before the avatar’s appearance. After a 1 ±0.1s interval, the avatar was first standing still for 1 ±0.1s at a virtual distance from the participant of 5m. The avatar then started moving towards the participant at a speed of 1.43 m/s. Participants were instructed to press the button as soon as they felt uncomfortable. In some trials, the avatar stopped as participants pressed the button, while at other trials (those with control success <100%), the button press did not always stop the avatar from approaching. If participants stopped the avatar successfully, the avatar remained still at its position until the next trial. The duration from the advent of the avatar until its disappearance was 3.5s.

**Fig. 1.**
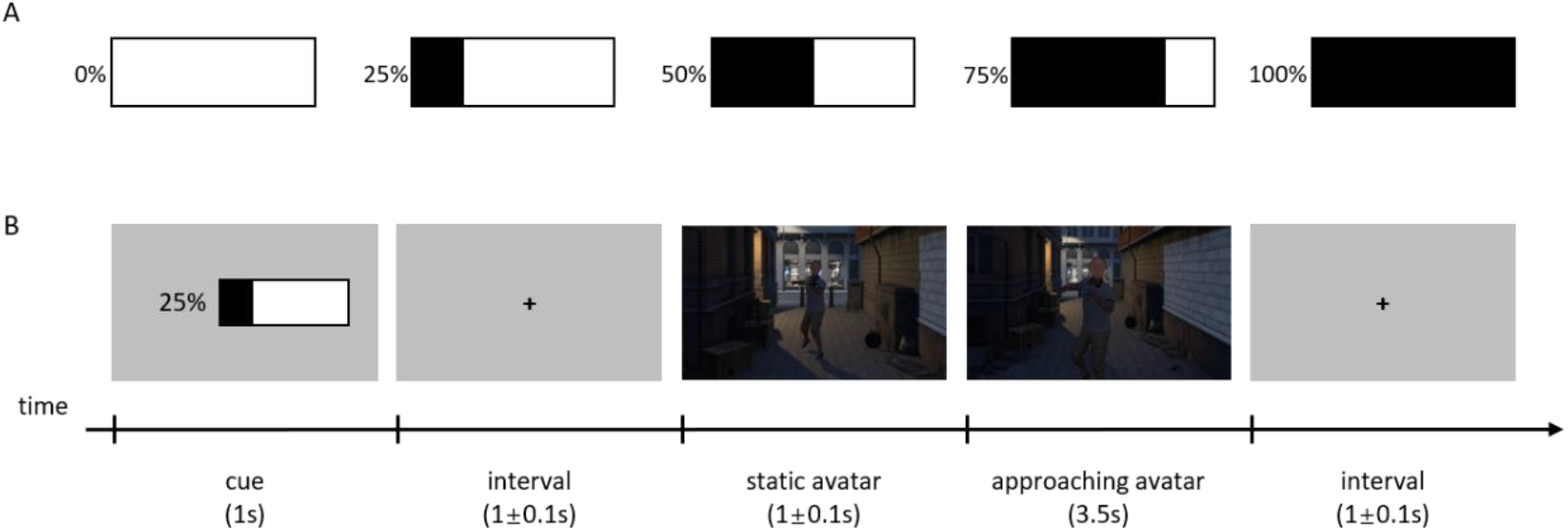
(A) Five kinds of controllable cues. (B) A trial procedure.

The study used a 5×2 within-subject design with five controllable cue conditions and two avatar emotions (angry, neutral) conditions. There were 40 trials per condition, and the total number of 400 trials was presented randomly in five runs, lasting in total 1h.

### 2.3 EEG acquisition

EEG data were recorded using an international 10-20 system, a scalp cap with 63 electrodes, and a sampling frequency of 250Hz (BrainVison Products, Munich, Germany). The electrode positioned on Cz was used as the reference during recording, and the forehead electrode positioned on FP1 was used as a ground electrode. Four electrodes were used to measure the electrooculogram (EOG). Two of them were used as vertical electrooculograms (VEOG). One was placed above the right eye, and another was placed below the right eye. The other two electrodes were used as a horizontal electrooculogram (HEOG), with one placed at the outer canthus of the left eye, and the other at the outer canthus of the right eye. Three electrodes were used for Electrocardiography (ECG). Two ECG electrodes were put one centimeter below the center of the left and right collarbones separately. The third ECG electrode was put on the right waist. The remaining 54 electrodes covered the whole scalp, including locations FPz, AFz, Fz, FCz, CPz, Pz, POz, Oz, AF7, AF8, AF3, AF4, F7, F8, F5, F6, F3, F4, F1, F2, FC5, FC6, FC3, FC4, FC1, FC2, T7, T8, C5, C6, C3, C4, C1, C2, TP7, TP8, CP5, CP6, CP3, CP4, CP1, CP2, P7, P8, P5, P6, P3, P4, P1, P2, PO7, PO8, PO3, PO4, O1, and O2. Impedances for reference and ground were maintained below 5kOhm and all other electrodes below 10kOhm. The heavy VR headset could potentially influence the EEG signal and cause head movement during the EEG data collection. To reduce these risks, we combined VR with a chin rest in our EEG experiment. After lowering impedance, the VR headset was carefully placed on the chin rest. The participants were standing against a height-adjustable bar chair in front of a high desk with the VR headset-Chin rest setup.

### 2.4 EEG data preprocessing

EEG data were preprocessed and analyzed using FieldTrip version 20220104 (Oostenveld et al., 2011) in Matlab R2021b (MathWorks, U.S.). The signal was first segmented into epochs from 1000ms pre-stimulus (the static avatar) to 2000ms post-stimulus and then filtered with a 0.1–30 Hz band-pass filter. EEG data at each electrode were re-referenced to the average of all electrodes. Artifact rejection was done using independent component analysis (ICA, logistic infomax ICA algorithm; Bell & Sejnowski, 1995); on average, 1.88 ± 0.33 (mean ± SD) components were removed per participant. Finally, single trials during which the peak amplitude exceeded 3 SD above/below the mean amplitude were rejected.

On average, 75.36% ± 5.27% (mean ± SD) trials were preserved and statistically analyzed per participant.

### 2.5 Event-related potential analyses

The ERP analyses performed were time locked to the presentation of the static avatar to derive clean ERPs in response to the still image. Here, a time window from 200ms before the onset of static avatar until 1000ms after the onset was extracted from each trial of the preprocessed data. A baseline correction was applied by subtracting the average amplitude during the interval (−200 ~ 0ms) before the onset of the static avatar. The segmented EEG for each participant was averaged for each experimental condition, resulting in ERPs used for further statistical analyses, which were performed using IBM SPSS Statistics 27 (IBM Corp., Armonk, NY, USA). In the ERP analysis, we focused on the ERPs elicited by social threatening/non-threatening body expressions (as represented by angry/neutral avatars) and their sensitivity to the level of threat control (controllable cues). We separated the EEG channels into five spatial clusters and identified for each region a prominent ERP component and centered a time window on its peak based on visual inspection of the overall ERP waveform, topographical distribution of grand-averaged ERP and previous studies (Chai et al., 2022; Cunningham et al., 2005; de Gelder et al., 2004; He et al., 2011; Luo et al., 2010; Stekelenburg & de Gelder, 2004; Van Heijnsbergen et al., 2007). The resulting ERP components and associated time windows are shown for each region in table 1. The mean amplitude was computed as the average of all electrodes within the cluster within the specific time window.

**Table 1.** Brain regions and electrodes of ERP components and associated time windows.

| ERPs | Brain regions | Electrodes | Time windows |
| --- | --- | --- | --- |
| N170 | Temporal | P7, P8, TP7, TP8, CP5, CP6, P5, P6 | 180-230ms |
| VPP | Central-occipital midline | Cz, CPz, Pz, POz, Oz | 200-250ms |
| P3 | Parietal | P5, P6, P7, P8, PO7, PO8 | 280-350ms |
| N3 | Frontal-central (with midline) | FCz, Cz, FC1, FC2, C1, C2 | 300-350ms |
| LPP | Frontal-central (without midline) | FC5, FC6, F5, F6, F7, F8, C5, C6 | 500-800ms |

A repeated-measures 5×2 ANOVA (Controllable cue: 0% / 25% / 50% / 75% / 100% × Avatar emotion: angry/neutral) was applied to the mean amplitudes; this was done for each ERP component separately. Degrees of freedom for F-ratios were corrected with the Greenhouse-Geisser method. Statistical differences were considered as significant given a *p* < .05. To control for type I errors, a Bonferroni correction was applied to the *p*-values associated with the main effects and interaction effects of every ERP component. Only corrected p-values were reported.

### 2.6 Time-frequency analyses

To assess temporal variations in oscillatory EEG power within the range from 1 −30Hz, we decomposed each trial using the complex Morlet wavelet transform (frequency-bin size: 1 Hz, three cycles per time window, time-bin size: 50ms). To reduce edge effects, we applied the time-frequency analysis to epochs of longer duration (corresponding to the duration of the preprocessed epochs before ERP computation; see above) and used a longer and earlier baseline in the interval (−500 ~ −100ms) before the onset of the static avatar. We focused the statistical analysis on oscillatory power in the theta (4-7Hz) band, based on literature showing that theta activity is related to the processing of threat and control, especially at frontal and central scalp regions (DeLaRosa et al., 2014; Lange et al., 2022; Ma et al., 2016). Thus electrodes positioned at Fz, FCz, Cz, F1, F2, FC1, FC2, C1, and C2 were selected for this analysis. Inspection of theta power revealed a peak between 100ms and 200ms after the onset time in the frontal central region consistently across conditions. Based on this observation, we extracted mean theta power during the time window (100 - 200ms) at the selected electrodes and statistically analyzed it using the same repeated measures ANOVAs as for the ERP analysis; see above.

### 2.7 ECG analyses

A time window from 500ms before static avatar onset to 4500ms after the onset was extracted from the continuous ECG data. The ECGdeli toolbox (Pilia et al., 2021) was used for analyzing heart rate. One participant’s ECG data was not recorded; thus, 25 participants’ ECG data were included in the analysis. The electrode that was placed under the left collarbone was selected for this analysis as it was positioned closest to the heart, giving the strongest signal. Statistical analyses were the same as for ERP and theta power; see above.

### 2.8 Behavioral analyses

We instructed participants to press the button as soon as they felt uncomfortable with the approaching avatar. In some trials, participants pressed the button once (4.81% trials), while in other trials, participants did not press the button or pressed it more than once (16.42% trials). Two behavioral indicators were recorded. First, the virtual distance between the participant and avatar at the time when participants first pressed the button. For this, the time at which the participants pressed the response button was multiplied by the speed by which the avatar was approaching and this was subsequently subtracted by the distance at which the avatar initially appeared (Distance = 3.5-Response Time*Speed). Second, the number of button presses was recorded. Like the physiological measures above, each behavioral indicator was subjected to a 5 (Controllable cue: 0% / 25% / 50% / 75% / 100%) × 2 (Avatar emotion: angry/neutral) repeated-measures ANOVA.

### 2.9 VR questionnaire

Information about the participants’ subjective experience during the VR scenario was obtained with a questionnaire, which participants filled in after the experiment (Seinfeld et al., 2016; Seinfeld et al., 2021). The individual questionnaire items are shown in Table 2.

**Table 2.**
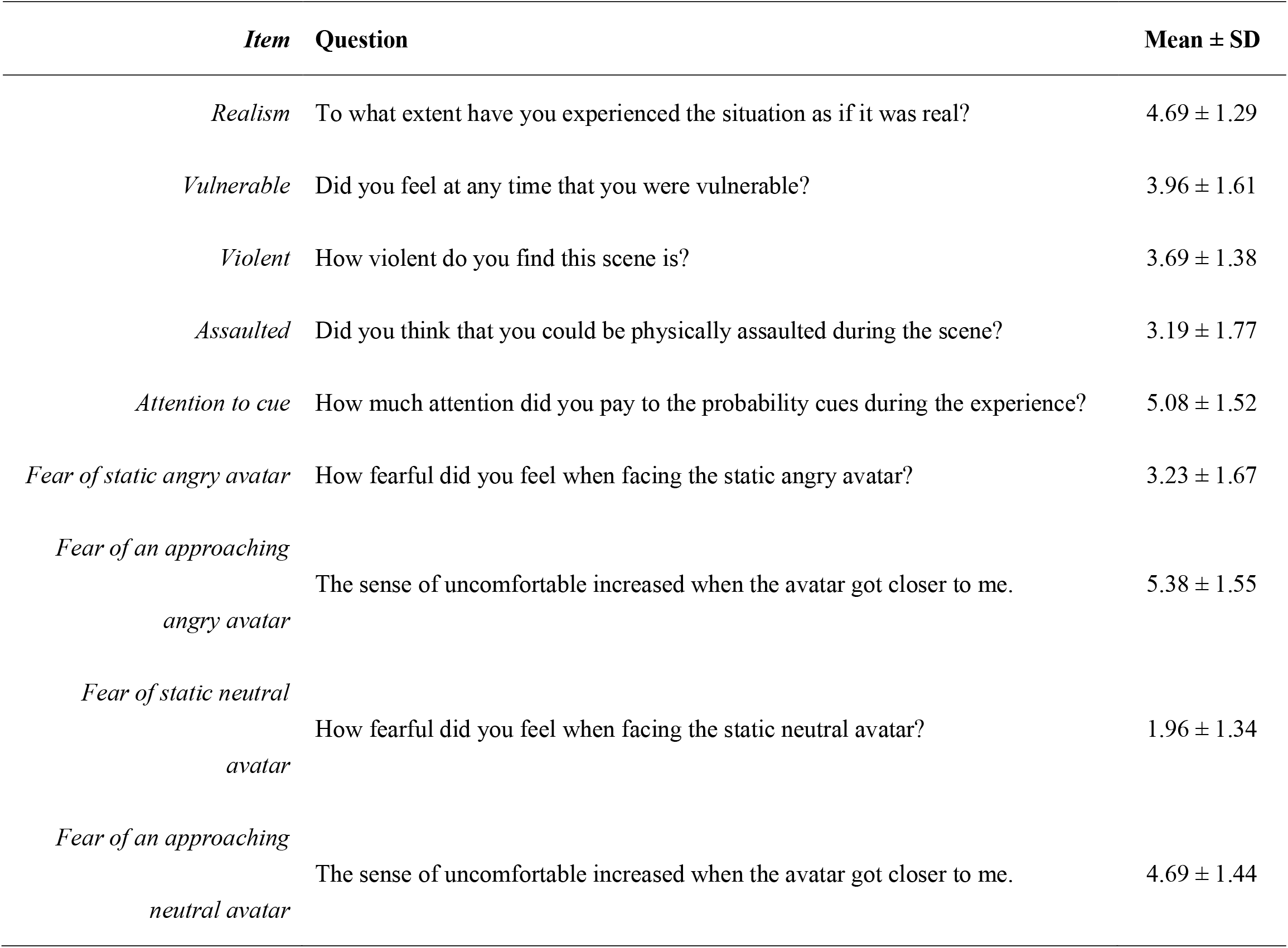
The items and mean ± SD rating scores in the VR questionnaire are shown. Ratings were made on a 7-point scale (1=not at all, 7=completely).

## 3. Results

### 3.1 VR questionnaire results

The items of the VR questionnaire and the mean ± SD of each item scores are shown in Table 2. We used a 7-point scale to test subjective feelings during the experiment, taking the median value “4” (the neutral subjective experience) as a reference to which we compared participants’ scores on each item. One-sample t-test results showed that realism (*t* (25) =2.74,*p* = .011), attention to cue (*t* (25) = 3.61, *p* = .001), fear of approaching angry avatar (*t* (25) = 4.55, *p* < .001) and fear of approaching neutral avatar (*t* (25) = 2.46, *p* < .021) were significantly larger than the reference value of 4, while assaulted (*t* (25) = −2.33, *p* = .028), fear of static angry avatar (*t* (25) = −2.36, *p* = .026), and fear of static neutral avatar (*t* (25) = −7.75, *p* < .001) were significantly smaller than the reference value. Furthermore, subjective experience of vulnerability (*t* (25) = −1.12, *p* = .904) and violence (*t* (25) = −1.14, *p* = .266) were not significant. Paired t-test results revealed that fear of a static angry avatar was significantly larger than fear of a static neutral avatar (*t* (25) = 4.44, *p* < .001), and fear of an approaching angry avatar was significantly larger than fear of an approaching neutral avatar (*t* (25) = 3.99,*p* < .001).

### 3.2 Behavioral results

For the first behavioral indicator (distance), the main effect of emotion was significant (*F* (1, 25) = 14.33,*p* < .001, *η_p_^2^* = 0.36) such that the distance between participants and the avatar was bigger when they saw the angry (2.24 ± 0.23) than neutral avatar (1.82 ± 0.21). The main effect of controllable cue was significant too (*F* (4, 100) = 4.56, *p* = .029, *η_p_^2^* = 0.15). A paired T-test between each controllable cue condition revealed no significant results after Bonferroni correction. The linear effect of controllable cue was significant (*F* (1, 25) = 5.07, *p* = .033, *η_p_^2^* = 0.17), showing that as the probability of the controllable cue increased, the tolerated distance decreased. The interaction effect between emotion and controllable cue was non-significant (*F* (4, 100) = 2.28, *p* = .096, *η_p_^2^* = 0.08) (Fig2A). Applying the same analyses to the second behavioral indicator (number of button presses) yielded qualitatively identical results, showing more button responses to the angry vs neutral avatar and to low vs. high probabilities of control success (main effect of avatar emotion: *F* (1, 25) = 4.43, *p* = .045, *η_p_^2^* = 0.15, angry: 1.61 ± 0.25, neutral avatar: 1.40 ± 0.18; main effect of controllable cue: (*F* (4, 100) = 6.15, *p* = .016, *η_p_^2^* = 0.20); linear effect of the controllable cue: *F* (1, 25) = 6.58, *p* = .017, *η_p_^2^* = 0.21; interaction effect (not significant): *F* (4, 100) = 1.83, *p* = .178, *η_p_^2^* = 0.07) (Fig2B).

**Fig. 2.**
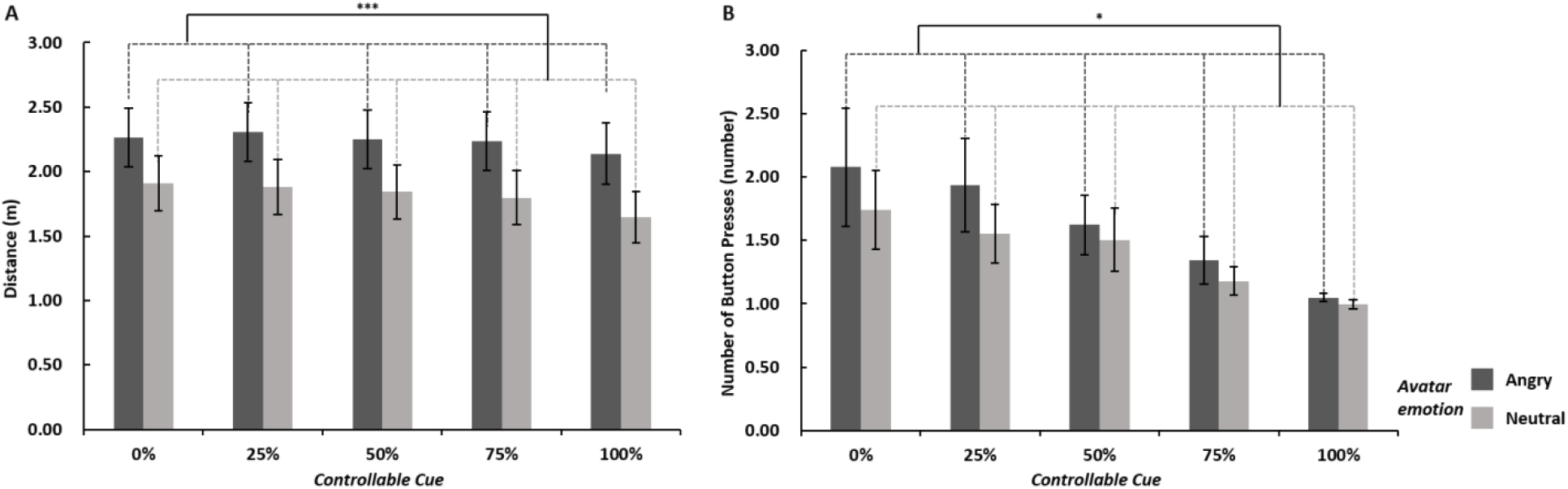
(A) Means and standard error (SE) of distance per condition. (B) Means and S.E. of the number of button presses per condition. \*\*\**p* < .001, *:*p* < .05

### 3.3 ERPs

#### 3.3.1 N170

The main effect of avatar emotion on the N170 amplitude was significant (*F* (1, 25) = 9.77, *p* = .017, *η_p_^2^* = 0.28) such that the angry avatar elicited larger N170 amplitudes (−2.04 ± 0.48 μV) than the neutral avatar (−1.68 ± 0.44 μV). The main effect of controllable cue (*F* (4, 100) = 0.34, *p* = 1, *η_p_^2^* = 0.01) and the interaction of the two factors were not significant (*F* (4, 100) = 1.23, *p* = 1, *η_p_^2^* = 0.05) (Fig3).

**Fig. 3.**
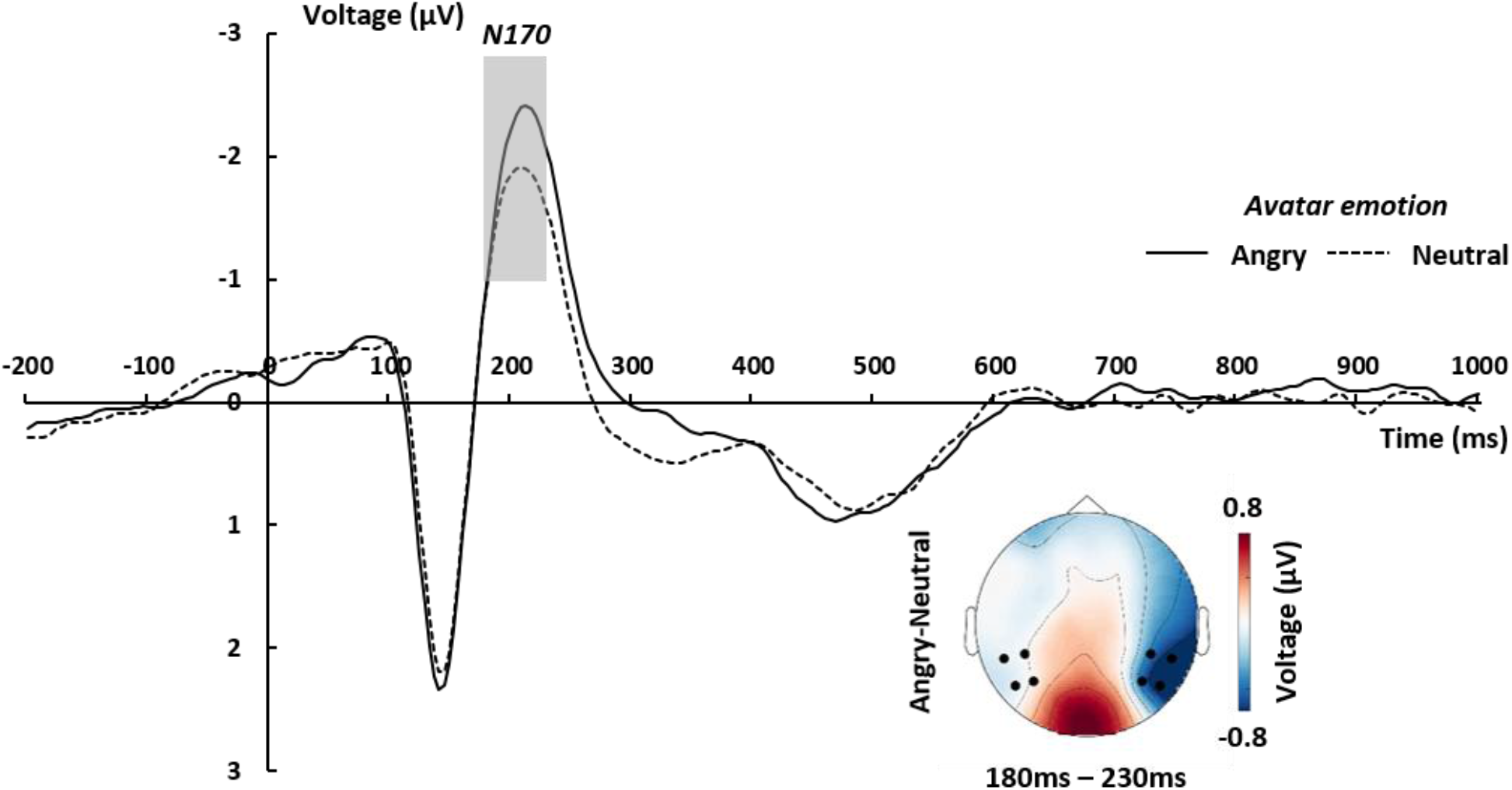
Grand-averaged ERP waveforms of N170 per avatar emotion condition. Waveforms were calculated by averaging the data at the electrodes P7, P8, TP7, TP8, CP5, CP6, P5, and P6, and across the controllable cue conditions. The “angry” minus “neutral” topographic map was calculated by averaging the data within a time window of 180 to 230ms after the onset of the static avatar. The black dots highlight the electrodes that were used to calculate grand-averaged ERPs.

#### 3.3.2 VPP

The main effect of avatar emotion on VPP was significant (*F* (1, 25) = 12.63, *p* = .006, *η_p_^2^* = 0.34) such that the angry avatar elicited larger VPP amplitudes (3.50 ± 0.53 μV) than the neutral avatar (2.93 ± 0.51 μV). The main effect of the controllable cue (*F* (4, 100) = 1.06, *p* = 1, *η_p_^2^* = 0.04) and the interaction of the two factors were not significant (*F* (4, 100) = 0.12, *p* = 1, *η_p_^2^* = 0.01) (Fig4).

**Fig. 4.**
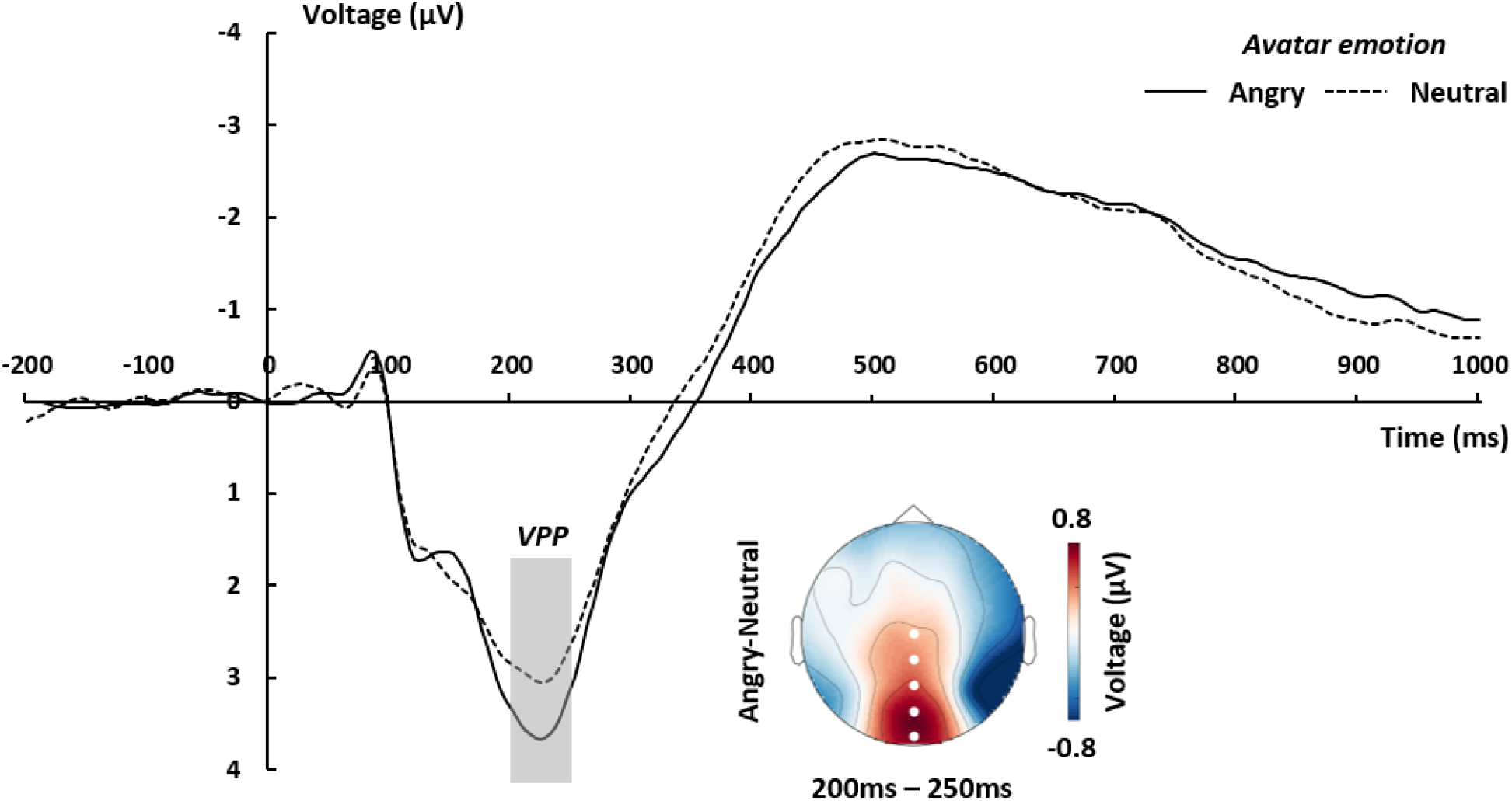
Grand-averaged ERP waveforms of VPP per avatar emotion condition. Waveforms were calculated by averaging the data at the electrodes Cz, CPz, Pz, POz, and Oz, and averaged across controllable cue conditions. The “angry” minus “neutral” topographic map was calculated by averaging the data within a time window of 200 to 250ms after the onset of the static avatar. The white dots highlight the electrodes that were used to calculate grand-averaged ERPs.

#### 3.3.3 N3

The main effect of avatar emotion on N3 was significant (F (1, 25) =9.28, *p* = .021, *η_p_^2^* = 0.27), such that the angry avatar elicited smaller amplitudes (−1.83 ± 0.25 μV) than the neutral (−2.12 ± 0.26 μV) avatar. The main effect of the controllable cue (*F* (4, 100) = 2.37, *p* = .300, *η_p_^2^* = 0.09) and the interaction of the two factors were not significant (*F* (4, 100) = 0.99, *p* = 1, *η_p_^2^* = 0.04) (Fig5).

**Fig. 5.**
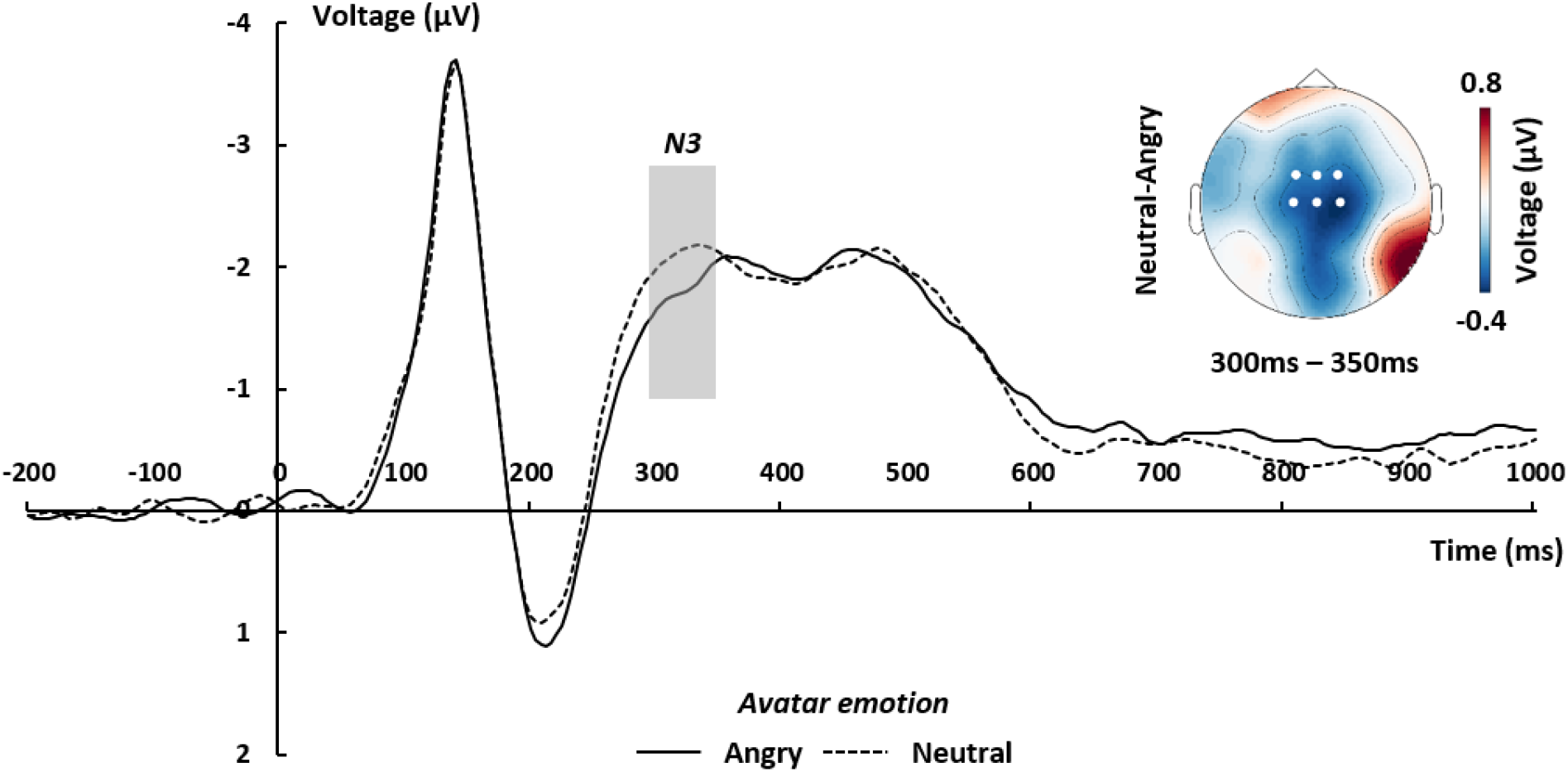
Grand-averaged ERP waveforms of frontal-central N3 per avatar emotion condition. Waveforms were calculated by averaging the data at the electrodes FCz, Cz, FC1, FC2, C1, and C2, and averaged across controllable cue conditions. The “neutral” minus “angry” topographic map was calculated by averaging the data within a time window of 300 to 350ms after the onset of the static avatar. The white dots highlight the electrodes which were used to calculate grand-averaged ERPs.

#### 3.3.4 P3

The main effect of avatar emotion on P3 was not significant (*F* (1, 25) = 4.81, *p* = .150, *η_p_^2^* = 0.16), such that the angry avatar elicited smaller amplitudes (2.43 ± 0.57 μV) than the neutral avatar (2.76 ± 0.57 μV). The main effect of controllable cue (*F* (4, 100) = 1.36, *p* = 1, *η_p_^2^* = 0.05) and the interaction of the two factors were not significant (*F* (4, 100) = 0.33, *p* = 1, *η_p_^2^* = 0.01).

#### 3.3.5 LPP

The main effect of controllable cue on LPP was significant (*F* (4, 100) = 4.24, *p* = .034, *η_p_^2^* = 0.15), such that the 100% cue elicited larger amplitudes (0.18 ± 0.25 μV) than the 75% cue (−2.56 ± 0.26 μV). The main effect of avatar emotion was not significant (*F* (1, 25) = 1.47, *p =* .950, *η_p_^2^* = 0.05). The interaction of controllable cue and avatar emotion was not significant (*F* (4, 100) = 2.68, *p* = .171, *η_p_^2^* = 0.10). (Fig6).

**Fig. 6.**
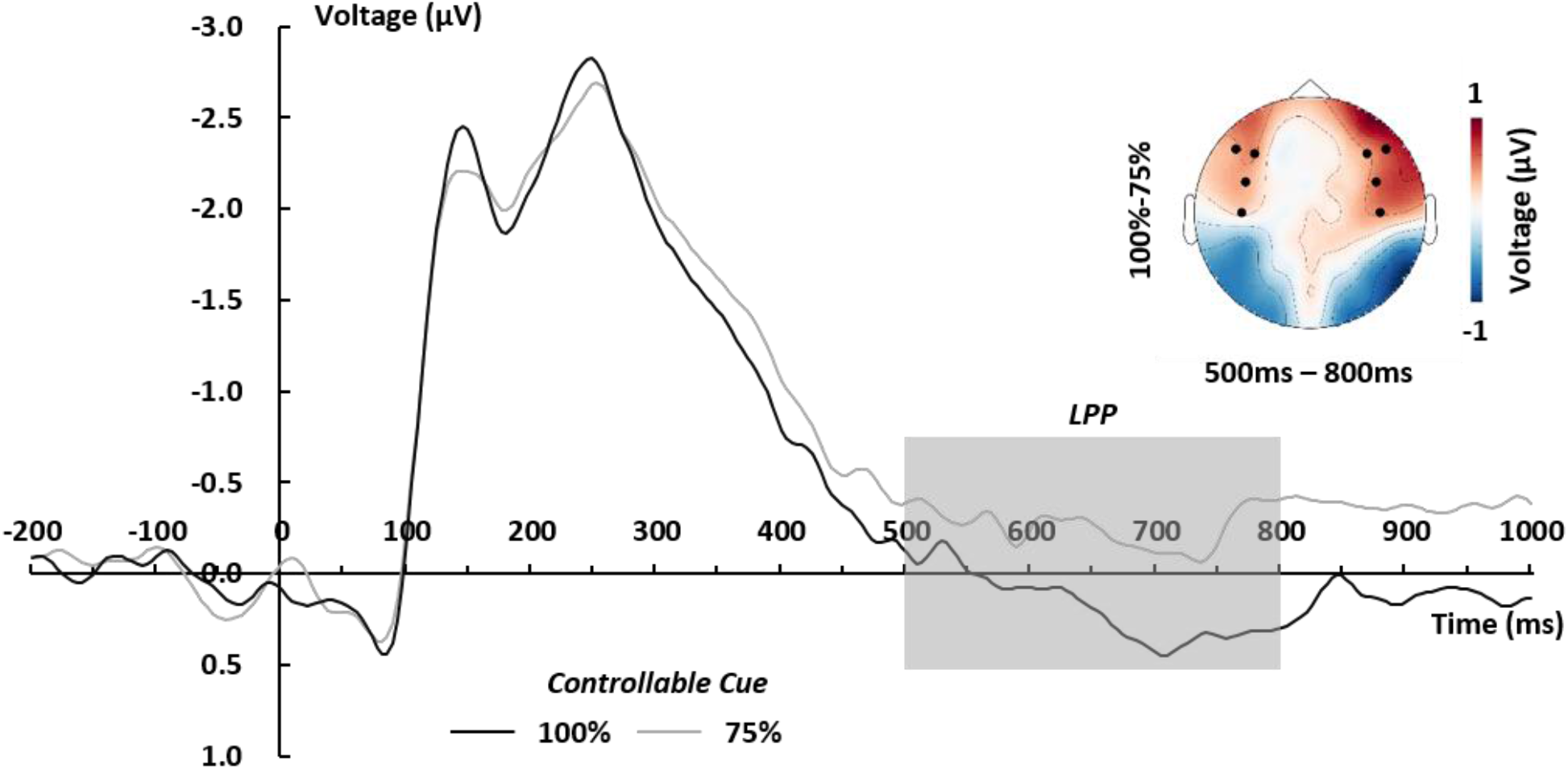
Grand-averaged ERP waveforms of frontal-center LPP under the 100% and 75% controllable cue conditions. Waveforms were calculated by averaging the data at the electrodes FC5, FC6, F5, F6, F7, F8, C5, and C6. The 100% controllable cue condition minus 75% controllable cue condition topographic map was calculated by averaging the data within a time window of 500 to 800ms after the onset of the static avatar. The black dots are highlighted electrodes which were used to calculate grand-averaged ERPs.

### 3.4 Time-frequency results

The main effect of avatar emotion on frontocentral theta power was significant (*F* (1, 25) = 7.87, *p* = .010, *η_p_^2^* = 0.24): theta power under the angry avatar condition (159.48 ± 20.68 dB) was increased compared to neutral avatar condition (140.24 ± 20.85 dB). The main effect of controllable cue (*F* (4, 100) = 0.84, *p* = .501, *η_p_^2^* = 0.03) and the interaction of the two factors were not significant (*F* (4, 100) = 0.74, *p* = .544, *η_p_^2^* = 0.03) (Fig7).

**Fig. 7.**
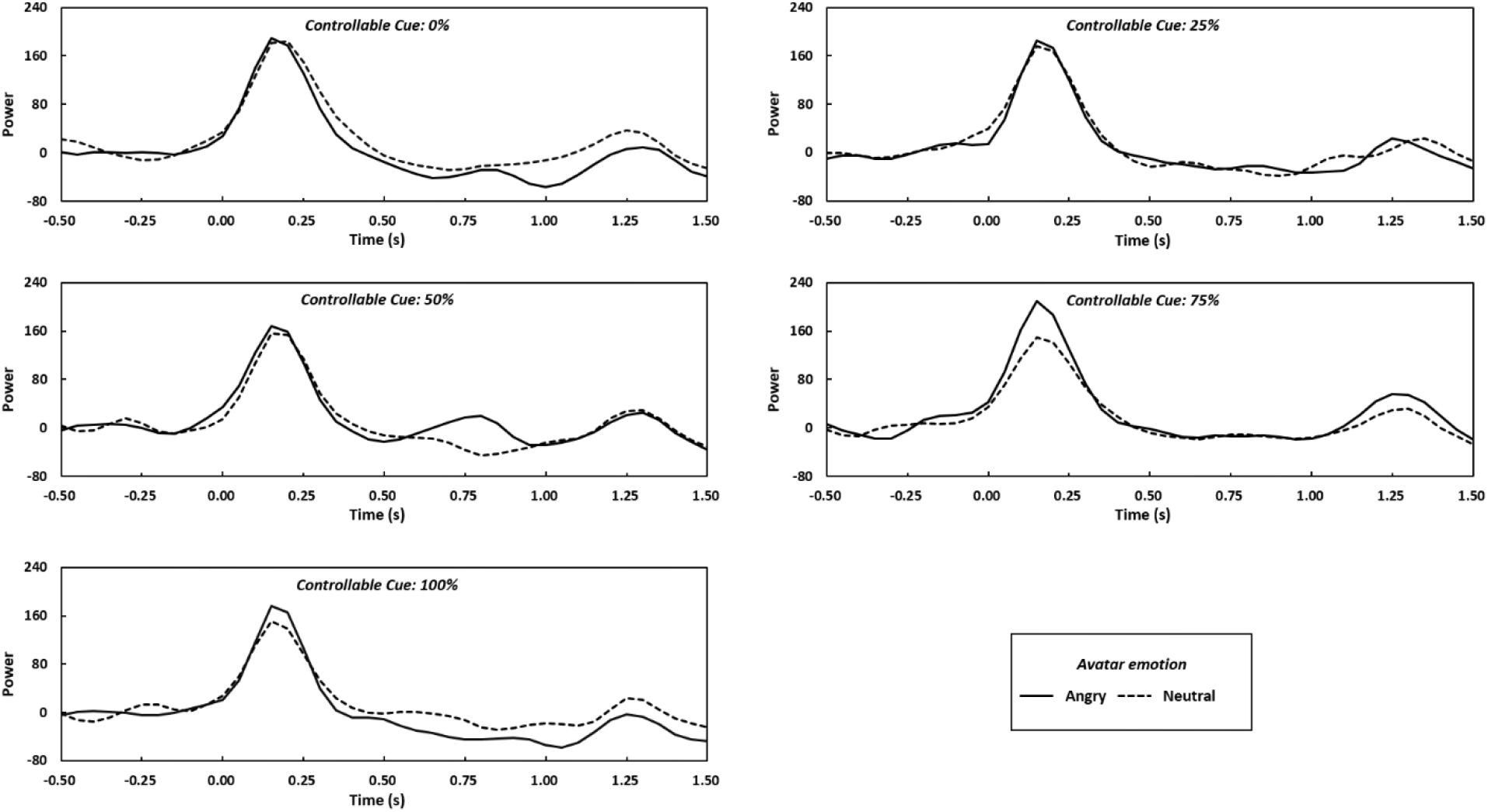
Theta power was calculated by averaging the theta band (4-7Hz) at the electrodes Fz, FCz, Cz, F1, F2, FC1, FC2, C1, and C2 per condition.

### 3.5 ECG results

Avatar emotion had a significant main effect on ECG (*F* (1, 24) = 7.482, *p* = .012, *η_p_^2^* = 0.24) such that the angry avatar elicited a higher heart rate (77.84 ± 2.49 beats per minute, BPM) than the neutral avatar (77.55 ± 2.45 BPM) did. The main effect of controllable cue (*F* (4, 96) = 1.24, *p* = .301, *η_p_^2^* = 0.05) and the interaction of the two factors were not significant (*F* (4, 96) = 0.45, *p* = .744, *η_p_^2^* = 0.02) (Fig8).

**Fig. 8.**
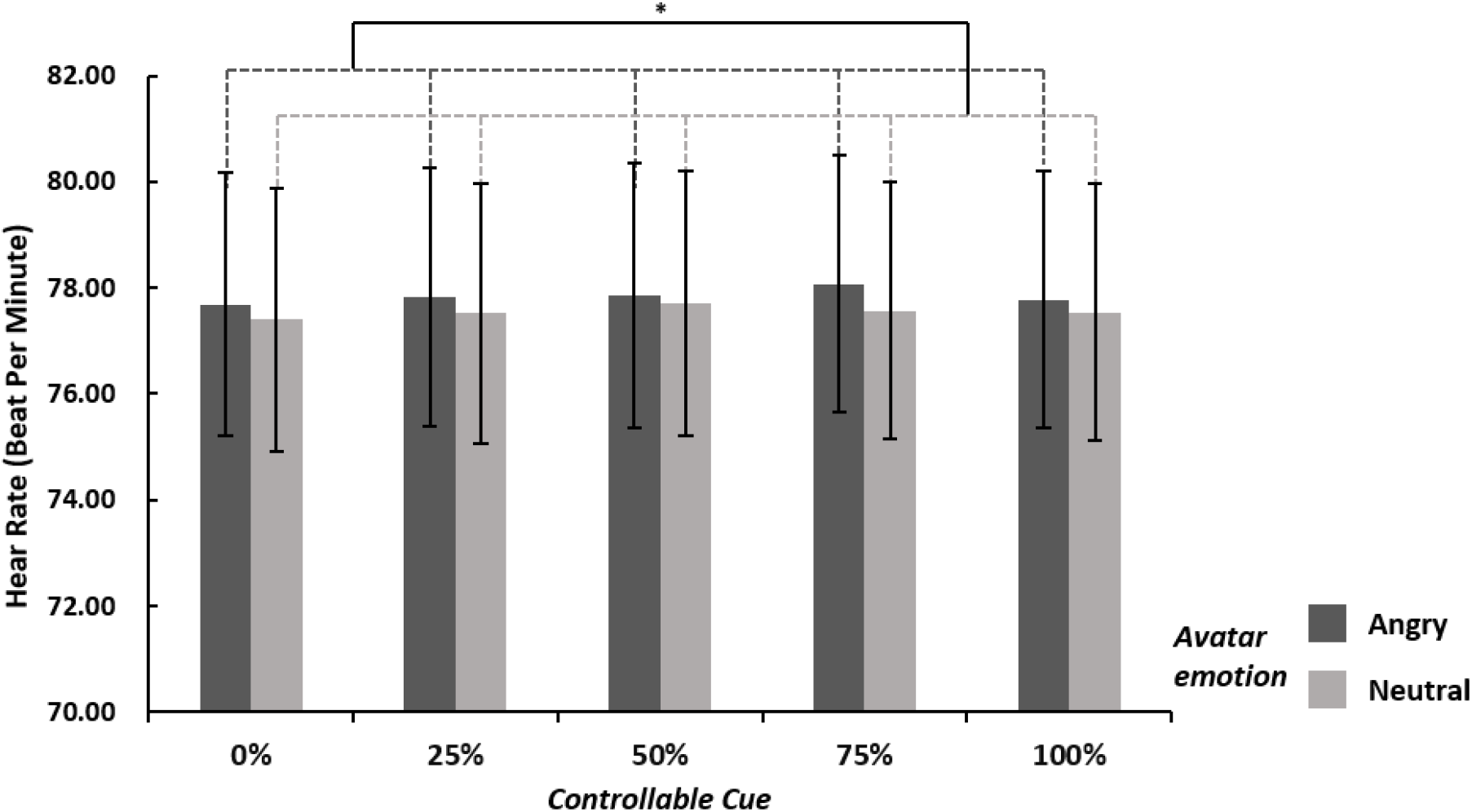
Means and SE of heart rate per condition. \**p* < .05

## 4. Discussion

In this study, we investigated the behavioral, EEG, and cardiac responses of human participants that were facing angry and neutral avatars in a VR environment in which they had control various degrees of control over the interaction with the avatar. Behaviorally, we observed a difference in the time/distance at which participants felt uncomfortable with the approaching avatar depending on the presence of threat. This is in line with the literature showing that threat imminence triggers defensive behavior (Blanchard & Blanchard, 1990; de Haan et al., 2016; Qi et al., 2018; Riem et al., 2019; Terburg et al., 2018). The impact of personal distance for social threat experience was first shown with full body expression of avatars in a study using VR and fMRI (de Borst et al., 2020). Combining VR with EEG in the present study now allows a detailed picture of the time courses. The questionnaire results also suggested that participants felt more threatened when facing an angry than a neutral avatar.

Concerning the impact of controllability, we found a significant effect of the controllable cue condition for each of the two behavioral indicators. First, as the probability of successful control increased, the distance from the avatar that participants judged tolerable decreased. This result is supported by a previous behavioral study (Iachini et al., 2016). Second, we observed that as the probability of successful control decreased, the number of button presses increased. This is consistent with the notion that the closer a threatening stimulus is to the self, the more likely the danger and the stronger the elicited defensive responses (Bufacchi, 2017). In our experiment, the button press was regarded as a defensive behavior. As the chances of successfully stopping the approaching avatar became higher, the number of button presses (defensive behavior) decreased.

At the neural level, we have three major findings. Seeing a threatening body expression (angry avatar) increased the amplitude of early ERP components (N170 and VPP) compared to non-threatening body expressions (neutral avatar). Furthermore, threatening body expressions elicited a smaller N3 than neutral body expressions. Finally, full control (100% controllable cue) increased the amplitude of the late component LPP as compared to the 75% controllable cue. Taken together, we show that social threat is detected in the early stages and independently of the possibility of control. In contrast, the impact of perceived control over the threat is reflected in the electrophysiological responses at later stages.

### Early threat detection

Our results indicate that participants were more sensitive to affective stimuli than neutral ones in the early stages of full body avatar processing. There are consistent but relatively few findings on body perception, a situation reflecting that whole body perception is still much less studied than face perception. Previous studies have reported that not only facial expressions but also whole body images trigger this early brain activity (Farzmahdi et al., 2021; Meeren et al., 2005; Stekelenburg & de Gelder, 2004; Van Heijnsbergen et al., 2007) and that the activity in this time window is sensitive to the emotional expression as shown by larger VPP amplitudes for a fearful than a neutral body (Stekelenburg & de Gelder, 2004). Also, consistent with our results, N170 and VPP seem to derive from a common source in the brain (Joyce & Rossion, 2005). An interesting finding consistent with the present results is that top down attention to the body stimulus did not influence the N170 amplitudes (Hietanen et al., 2014), indicating that body expression perception is an automatic stimulus driven process. Here we add to this by showing that clear knowledge of subjective control of the threat does not impact the course of early body expression perception. This result suggests that we are observing here the early stages of threat perception, that are then followed by calculations of alternative escape decisions (Qi et al., 2018). Given this interpretation of the processes associated with N170, it is worth stressing that our results were obtained in a VR setting which is characterized by an immersive experience of realism but also at the same time, a subjective understanding that the experience is not real, in our case that the participant is not really threatened. An alternative outcome might have been that participants knowledge of the danger being ‘unreal’ would have overruled this early signature of threat experience.

### Temporal dynamics of behavioral control

The middle-late component N3 is related to source allocation and response preparation. A lower amplitude of the N3 component is thought to reflect that more cognitive resources and brain resources are being mobilized to prepare for a response to the threat (Coenen, 1995; Ke et al., 2022; Mayer et al., 2021). E.g., higher cognitive tasks have been shown to elicit lower N3 amplitudes than simpler tasks (Michida et al., 1998). Moreover, negative emotional visual stimuli have been observed to evoke lower N3 than positive emotional ones (Ke et al., 2022). In our experiment, we found threatening body expressions to elicit lower N3 than neutral body expressions, suggesting that the threatening stimuli elicited negative emotions and required more cognitive resources than the neutral stimuli.

Concerning the late positive potential (LPP), previous studies reported that LPP refers to task-relevant, motivational engagement and action preparation during the later stage (Di Lemma et al., 2020; Gable et al., 2015; Gantiva et al., 2020). Johnen and Harrison found that LPP amplitude was larger under conditions of certainty compared to less certain conditions (Johnen & Harrison, 2020). In line with this study, we found that perfect control (100% controllable cue) elicited larger LPP than the 75% controllable cue. This suggests that perfect control opportunity resulted in more motivation engagement than 75% control success.

We also examined oscillatory brain responses in relation to differences in successful control probability for threatening and neutral body expression. Our data show that theta power in frontal central regions was only modulated by avatar emotion. There was a significant increase in response to the threatening body expression (angry avatar) compared with the neutral body expression (neutral avatar). This is consistent with findings showing that increased theta power is related to higher emotional arousal (Aftanas et al., 2001; Aftanas et al., 2002; Sulpizio et al., 2021) and that greater theta power may be induced by social threat compared with non-threat stimuli (Diao et al., 2017).

Our ECG results are consistent with the literature showing a higher heart rate for social threat than for non-threat (Eisenbarth et al., 2016; Weeks & Zoccola, 2015), although another study found a lower heart rate for threatening vs neutral body expressions (Mello et al., 2022). In that study, participants passively viewed a threatening avatar coming closer, which induced the freezing response reflected in a reduced heart rate. In our paradigm, participants could actively stop the approaching avatar from coming closer by pressing the button. Thus, the increased heart rate might reflect emotional arousal to the threatening body expression rather than a freezing response.

## 5. Conclusion

The amplitudes of the earlier components (N170/VPP/N3) are elicited by viewing a threatening body expression and seem to be independent of control opportunities, while the latter modulate the later LPP component. Our findings on N170/VPP effects show that these two components may be modulated by threatening/neutral body expressions, which may reflect mechanisms involved in rapid detection of social threat in an early-middle stage, such as decoding the meaning of a threatening body expression. The social threat is further processed in later stages, as indicated by the effects of avatar emotion on the middle-late cognitive components N3. The ability to control the threat shows in the late cognitive evaluation stages, as reflected by the LPP effects. In addition, the increased frontal central theta power and heart rate are related to social threat processing. In sum, our study provides behavioral and neural insights into how humans process social threats under varying levels of control. On the methodological side, our study presents a novel VR-EEG-ECG setup that will be useful also for future VR and EEG studies investigating social interaction situations in a naturalistic fashion.

## Acknowledgement

This work was supported by the European Research Council (ERC) FP7-IDEAS-ERC (Grant agreement number 295673; Emobodies), by the ERC Synergy grant (Grant agreement 856495; Relevance), by the Future and Emerging Technologies (FET) Proactive Program H2020-EU.1.2.2 (Grant agreement 824160; EnTimeMent), by the Industrial Leadership Program H2020-EU.1.2.2 (Grant agreement 825079; MindSpaces), by the Horizon 2020 Programme H2020-FETPROACT-2020-2 (grant 101017884 GuestXR) and by China Scholarship Council (CSC202008440538).

## Data availability

The data are available upon request from the corresponding author.

